# Long live the host! Proteomic analysis reveals possible strategies for parasitic manipulation of its social host

**DOI:** 10.1101/2022.12.23.521666

**Authors:** Juliane Hartke, Alejandro Ceron-Noriega, Marah Stoldt, Tom Sistermans, Marion Kever, Jenny Fuchs, Falk Butter, Susanne Foitzik

## Abstract

Parasites with complex lifecycles often manipulate the phenotype of their intermediate hosts to increase the probability of transmission to their definitive hosts. Infection with *Anomotaenia brevis*, a cestode that uses *Temnothorax nylanderi* ants as intermediate hosts, leads to a multiple-fold extension of host lifespan and to changes in behaviour, morphology, and colouration. The mechanisms behind these changes are unknown, as is whether the increased longevity is achieved through parasite manipulation. Here we demonstrate that the parasite releases proteins into its host with functions that might explain the observed changes. These parasitic proteins make up a substantial portion of the proteome of the hosts’ haemolymph, and thioredoxin peroxidase and superoxide dismutase, two antioxidants, exhibited the highest abundances among them. The largest part of the secreted proteins could not be annotated, indicating they are either novel or severely altered during recent coevolution to function in host manipulation. We also detected shifts in the hosts’ proteome with infection, in particular an overabundance of vitellogenin-like-A in infected ants, a protein that regulates division of labour in *Temnothorax* ants, which could explain the observed behavioural changes. Our results thus point at two different strategies likely employed by this parasite to manipulate its host – by secretion of proteins with immediate influence on the host’s phenotype and by altering the host’s translational activity. Our findings reveal the intricate molecular interplay required to influence the phenotype of a host and shed light on potential signalling pathways and genes involved in parasite-host communication.

## Introduction

The closest molecular interactions and antagonistic dynamics occur between parasites and their hosts, resulting in highly specialized morphologies, physiologies, and life cycles of parasites that can be characterized as either direct or indirect (Olsen, 1986). Parasites with a direct life cycle transmit themselves or their progeny directly from one host to the next. In contrast, parasites with an indirect or ‘complex’ life cycle require at least one intermediate host or vector before they can complete their life cycle in the definitive host (Olsen, 1986). The common denominator of parasites with complex life cycles is that they require a host switch to complete their life cycle. Though difficult to verify experimentally, many of these parasites are thought to actively increase the likelihood of transmission. The malaria parasite *Plasmodium falciparum* alters the odour of their human hosts, possibly to increase their attractiveness towards mosquitoes during the infective stage (de Moraes et al., 2014; Lacroix et al., 2005). *Toxoplasma gondii* increases the risk-taking behaviour of infected rats and causes them to be attracted to the smell of cat urine, likely facilitating predation by the definitive host (Berdoy et al., 2000; Frenkel et al., 1970; Golcu et al., 2014). Those direct changes of host behaviour or odour are examples of adaptive host manipulation by the parasite. Despite the description of numerous case studies in which host manipulation appears to occur, we lack general knowledge of the underlying mechanisms by which parasites alter the phenotype or behaviour of their host. Amongst others, proposed pathways of manipulation are via energy deprivation (Lafferty & Shaw, 2013), via the parasites location within the body of the host (e.g., nervous system, or around muscle), and via the release of manipulative agents (Martin et al., 2015). The difficulty in defining the exact mechanism by which a change in host phenotype is achieved arises from the distinction between parasite-induced effects and host defence mechanisms, but also from the fact that often several interconnected signalling pathways are affected during an infection (Kavaliers et al., 1999). One particularly interesting parasite for which increased transmission due to host manipulation has been postulated (Beros et al., 2015), is the cestode *Anomotaenia brevis*. This parasite has an indirect life cycle with the ant *Temnothorax nylanderi* as an intermediate host and two species of woodpecker (*Dendrocopos major* and *D. minor*) as definitive hosts (Plateaux, 1972). The infection of the intermediate host happens when foraging ants bring bird faeces with cestode eggs into the nest and feed those to the developing larvae. Within the ant, the cestode eggs hatch into larvae and pass through the gut wall into the haemocoel, where they develop into cysticercoids. When a woodpecker opens acorns or sticks, in which *T. nylanderi* build their nests, and feeds on the infected ants, the cysticercoids develop into adult tapeworms and complete their life cycle (Figure 1).

**Fig. 1.**
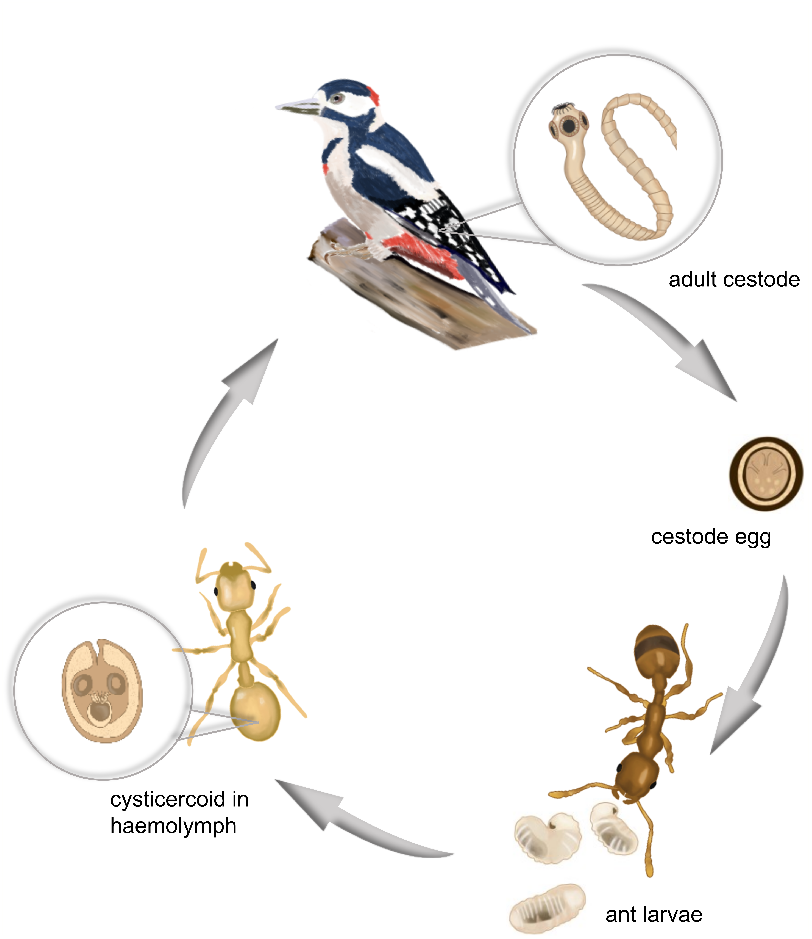
Lifecycle of *Anomotaenia brevis*. The adult stages in different woodpecker species are producing eggs that are excreted by the bird and subsequently fed to growing ant larvae of *Temnothorax nylanderi*. The eggs develop into cysticercoids that are attached to the gut in the hemocoel. The lifecycle is completed when infected ants are eaten by the woodpecker.

Infected ants have a yellow, less pigmented and less sclerotized cuticle, in contrast to the brown colour of healthy workers (Plateaux, 1972; see Figure 1). They are less active than their non-infected nest mates and rarely leave the nest but are fed and groomed more frequently (Feldmeyer et al., 2016; Scharf et al., 2012). The most interesting change, however, is in their lifespan. Over a three-year period, infected workers showed no difference in survival compared to queens, which can live up to two decades in this species (Plateaux, 1986). In contrast, all uninfected, even newly emerged workers died during this time (Beros et al., 2021). Longer-term observations that could provide information on the maximum life expectancy of infected individuals are lacking so far, but anecdotal observations show that infected workers can live up to seven years (A. Buschinger, pers. Obs.). Both the extended lifespan and the fact that parasitized individuals remain in the nest even when disturbed (Beros et al., 2015) are factors that increase the likelihood of transmission to the final host and may be the result of active manipulation. In this study, we aimed to further unravel the target mechanisms for the observed increase in lifespan of infected individuals. Previous studies investigating the factors contributing to the longer-lived phenotype of infected ants focused mainly on changes in gene expression after infection (Feldmeyer et al., 2016; Stoldt et al., 2021). Another promising target to identify which molecular pathways are affected by an infection, however, is the proteome. The proteome reflects the biological activity and downstream effects of infection even more directly and includes not only the proteins synthesized by the host itself, but possibly also those released by the parasite through active secretion or passive excretion. By analysing the proteome of the haemolymph, it is possible to identify circulating substances, whether parasitic or host in origin, that increase the fitness of the parasite. To be able to detect active secretion of proteins by the cestode, we analysed the haemolymph of infected and uninfected workers of *T. nylanderi* for the presence of secreted proteins of the cestode *A. brevis*. In addition, we compared the abundance of proteins from whole-parasite proteomes to those found in the haemolymph, as this would provide further evidence of active delivery of some proteins into the host organism and thus possibly of host manipulation. We also compared workers and queens from infected and uninfected colonies, which gives insight into the consequences of infection for the nestmates of infected ants, which also show an altered phenotype, e.g., increased mortality. Finally, a caste comparison was of interest, as queens show a similar low mortality rate as infected workers, a similarity that could have the same or a different molecular basis. Although we worked with very limited material, as *Temnothorax* ants and their colonies are very small and infected individuals are rare, we were able to (*i*) identify proteins with potentially manipulative functions (i.e. the cestode proteins) and (*ii*) uncover changes in the ant proteome that may be the result of the secreted parasite proteins.

## Methods

### Ant collection and maintenance

*Temnothorax nylanderi* colonies were collected in the Lenneberg Forest in Mainz (50°00’41.7”N; 8°10’32.8”E). In the laboratory, the colonies were moved to a small slide nest covered with red foil to give them a darkened environment, which was placed in a larger Plexiglas nest, consisting of three chambers, with a plastered floor. They received water and honey and were kept in the laboratory at constant conditions of 20°C and 70% humidity for an acclimation period of seven days before haemolymph extractions.

### Haemolymph and cestode sampling

Haemolymph was extracted for proteomic analyses from three different groups of ants: from infected worker ants, from non-infected worker ants that belong to the same colony (hereafter short: nestmate ants), and non-infected worker ants from healthy colonies (hereafter short: healthy ants). For each extraction, the haemolymph of ten individual ants were pooled, and we performed four biological replicates. For each colony that was included in the proteome analysis, we also extracted the haemolymph of the (uninfected) queen from the colony, which resulted in the same four replicates for both infected and healthy colonies, but with only one individual each.

For the collection of the haemolymph, the ants were immobilized by placing them head first into a flexible foam plug, so that the abdomen was freely accessible. By creating a small perforation with a micro scissor between two segments of the abdomen, we gained access to the haemolymph that was collected with a narrowed capillary (Hirschmann, Eberstadt, Germany) to ensure that no cestodes were transferred. The collected clear fluid was transferred into a 1.5 ml reaction tube filled with 8 M urea (Sigma Aldrich). All samples were stored at -20°C until mass spectrometry analysis.

Following haemolymph extraction, the gaster of each individual was dissected to confirm the infection status of the individual and to collect the cysticercoid cestodes for proteomic analyses. Cysticercoids reside in the haemolymph in close vicinity to the midgut, sometimes loosely attached to it. Individual worker ants can contain between 1 to 73 cestodes (T. Sistermans, unpublished). We detected between 1 to 36 parasites per individual and our four replicates contained a total of 126, 45, 100 and 67 cysticercoids.

### Mass spectrometry analysis

The haemolymph solution was diluted with 150 μl 50 mM ammonium bicarbonate (ABC) buffer pH 8.0. The samples (ca. 200 μl) were reduced with 10 mM f.c. dithiotreitol (Sigma Aldrich) for 40 min and subsequently alkylated in the dark with 50 mM f.c. iodoacetamide (Sigma Aldrich) for 40 min. Afterwards 500 ng LysC protease (Wako) was added and after 3 hours of incubation the samples were supplemented with 500 ng MS-grade trypsin (Serva) and digested overnight. The peptides were loaded onto a C18 StageTip (Rappsilber et al., 2007) and stored until measurement. The eluted peptides were separated on a heated 50-cm reverse-phase capillary (75 μM inner diameter) packed in-house with Reprosil C18 material (Dr. Maisch GmbH). Peptides were eluted along an optimized 90 min gradient from 6 to 40% Mixture B (80% ACN/0.5% formic acid) with an EASY-nLC 1,200 system (Thermo Fisher Scientific). Spray voltage was set to 2.2 kV. Measurement was done on an Orbitrap Exploris 480 mass spectrometer (Thermo Fisher Scientific) operated with a Top20 data-dependent MS/MS acquisition method per full scan. The full scan had a resolution of 60,000 with a scan window set from 350-1,600 m/z. The isolation window was 1.4 m/z. The MS/MS fragment scan was obtained with a resolution of 15,000 after HCD fragmentation. Fragmentation was restricted to peptides with charge state 2-6. All raw files were processed with MaxQuant version 1.6.0.5 (Cox & Mann, 2008) and peptides were matched to the protein database of *T. nylanderi* with 43,252 entries (Stoldt et al. 2021) and the three-frame translated (at least 100 aa ORFs) protein sequences of the cestode *A. brevis* with 173,911 entries (Stoldt et al., 2021). MaxQuant was used with pre-set standard settings, but we activated the following options: match between runs, label-free quantitation (LFQ) and intensity based absolute quantification (iBAQ), and deactivated fastLFQ. LFQ quantitation was only performed on unique peptides. Prior to further analysis, contaminants, reverse hits and proteins that were only identified by a modified peptide, were removed. The statistical analyses were done with R Studio (v3.6.2).

### Analyses of host proteins

We employed two separate bioinformatic approaches to analyse the *T. nylanderi* and *A. brevis* proteins. For *T. nylanderi* proteins, abundances of the different groups (healthy, nestmates, and infected) were compared in a pair-wise fashion using a Welch t-test and the mean difference of log2 transformed and imputed LFQ intensities between the different conditions were computed. This was repeated for a comparison of queens from healthy colonies and queens from infected colonies. To test whether infected workers show higher abundances of proteins that are usually only found in queens, we compared proteins that are significantly more abundant in queens (compared to healthy and nestmate workers), to those that are significantly more abundant in infected workers (compared to their nestmates and to healthy workers). Since *T. nylanderi* is usually monogynous, we could only use one queen per nest for the mass spectrometry analysis, as opposed to 10 pooled workers, rendering a direct comparison of iBAQ between our queen and worker samples impossible. Thus, for pairwise comparisons between queens and workers, we first filtered out proteins that were only present in one dataset and then calculated the relative protein abundances (iBAQ single protein / iBAQ sum of all proteins) separately for each of the four replicates. The pairwise comparisons were done in the same way as described above.

In addition, we extracted proteins that were unique to nestmate queens (queens from infected colonies) compared to nestmate workers, and in healthy queens compared to healthy workers. We checked whether those proteins tended to occur in higher abundances in infected workers, both compared to their nestmates and healthy workers, and whether also infected workers express those proteins uniquely.

Putative protein functions were obtained by performing a BLASTp search against the non-redundant invertebrate protein database (accessed: 14.09.2021). Proteins were functionally annotated using InterProScan v5.46-81.0. A Gene Ontology (GO) term enrichment analysis of biological processes was conducted with topGO (Alexa & Rahnenführer, 2016) compared to the *T. nylanderi* transcriptome. We used the weight01 algorithm and checked for significance using Fisher statistics with a p-value cut-off of p=0.05. Furthermore, a text mining approach was conducted to assign functional descriptions from reviewed entries of the UniProt database for *Drosophila* to the identified proteins. We then searched those functions for terms that are associated with longevity or ageing, immunity, transport, stress, and epigenetics, as we expected those functions to be of particular interest as a target for host manipulation, (for list of search terms see Supplement 1 Table S9) to identify potential candidate genes implicated in infection phenotypes.

During the analysis of the host proteome, we found vitellogenin (Vg) proteins in differential abundances between the groups. Vgs have diversified in ants and are known to play a role in caste differentiation. To identify the type of Vgs, we aligned the protein sequences to the protein sequences of different insect Vgs obtained by Kohlmeier et al. (2018) with MAFFT (v7.487; Madeira et al., 2019) using standard settings. The resulting tree (see Supplement 1 Figure S8) was visualized using the iTOL online tool (Letunic & Bork, 2019).

### Analysis of parasite proteins

A similar approach was employed for the cestode proteins. Here, no differential abundance analysis was conducted, since cestode proteins should only be found in infected ants, thus rendering comparative analyses impossible. We therefore conducted all following analyses with proteins that were (i) found in at least two out of four replicates, and (ii) with proteins that were present in all four replicates. With those proteins, as for the ant proteins, we conducted a BLASTp and Interproscan search to obtain functional characterizations and GO terms. We searched reviewed UniProt entries of *C. elegans* to assign functions to the identified proteins. [la-belfont=bf]caption Those lists were then also searched for terms that are associated with longevity or ageing, immunity, transport, stress and epigenetics (Supplement 1 Table S9). The functional annotation of cestode proteins, by only using BLASTp, proved to be more difficult than for the ant host, since not many cestode genomes have been annotated, and the closest relative with an annotated genome is *Echinococcus granulosus*. We thus additionally chose those 15 cestode proteins with the highest abundances in the haemolymph of infected ants and employed a eukaryotic feature-based function prediction with FFPred 3 (Cozzetto et al., 2016). We furthermore identified significantly enriched GO terms with the use of topGO as described above (Alexa & Rahnenführer, 2016) in a comparison to the transcriptome of *A. brevis*.

To identify whether cestode proteins were found in the haemolymph of ants due to secretion, we ran SignalP 5.0 (Nielsen et al., 2019) on all identified cestode proteins, to predict the presence of signal peptides, as well as SecretomeP 1.0f (Bendtsen et al., 2004), to identify non-classical (i.e., not signal peptide triggered) protein secretion. For the SecretomeP analyses, all proteins were considered as putatively secreted when they showed a neural network score > 0.6, according to Bendtsen et al. (2004). We calculated the share of proteins with SignalP sites for the whole known *A. brevis* protein set as well as for all proteins present in at least one replicate, in at least two replicates, in at least three replicates, and in all four replicates. We compared the share of proteins with SignalP sites among those groups using a χ^2^ test.

For the identification of cestode candidate proteins that might contribute to the phenotypic changes observed in infected ants, we used only those proteins that are putatively secreted, were annotated in other organisms, have functions in longevity or immunity, and were found in at least two / all four replicates.

As the resulting protein abundances of whole cestode mass spectrometry analysis and haemolymph analysis are not directly comparable, we employed the same approach as for the comparison of ant proteins between queens and workers. Briefly, we filtered out proteins that were only found in one of the measurements of haemolymph and whole cestode samples and calculated relative abundances of the remaining proteins. Those values were used for a Welch t-test. Significantly more abundant proteins of either haemolymph or whole cestode samples were identified based on log10 transformed p-values and log2fold change differences of the relative abundances. We furthermore compared the percentage of annotated proteins between haemolymph and whole cestode data, and between proteins which are predicted to be secreted and non-secreted. We compared the two datasets also regarding the results of automated word category searches for longevity, immunity, transport, and epigenetics.

## Results

We identified 263 proteins from the haemolymph of infected ants that matched the transcriptome of *A. brevis* and that were present in at least two of four biological replicates. It should be noted that an identification of protein functions proved difficult. Only 137 proteins could be functionally identified via a BLAST search. For the remaining 126 proteins, either no BLAST result was found (112 proteins), or the proteins from related species were annotated as “hypothetical protein” (14 proteins).

We compared the number of proteins which were putatively secreted between proteins that were found in at least one (two, three or all four) biological replicate (Supplement 1 Figure S1). While the percentage of putatively secreted proteins vs non-secreted proteins was constant across the number of replicates, in which proteins were found, the share of proteins with predicted signal peptide cleavage (SignalP) sites, indicating their secretion, increased with the number of replicates (Figure 2 D). Compared to the percentage of proteins with SignalP sites that were predicted across the complete proteome of *A. brevis* (8.6%), we found a higher-than-expected proportion of those proteins present in the haemolymph with this molecular indicator of secretion (present in at least one replicate (20.3%): χ^2^ = 5, p = 0.03; present in at least two replicates (20.5%): χ^2^ = 5, p = 0.03; present in at least three replicates (28.3%): χ^2^ = 12, p = 0.0007; present in all four replicates (33.7%): χ^2^ = 17, p = 0. 00003).

**Fig. 2.**
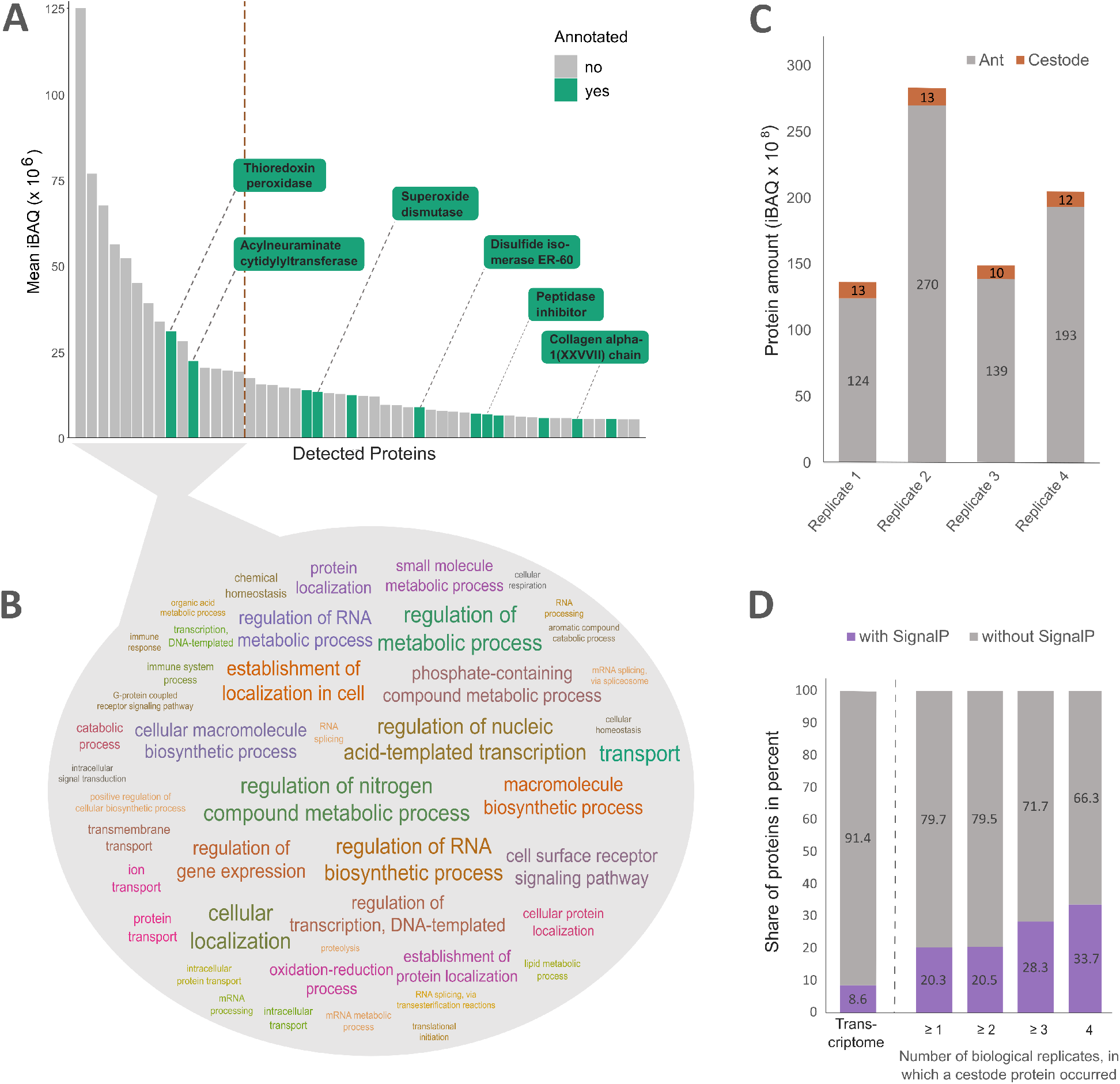
A) Mean sum of the intensity based absolute quantification (iBAQ) of peptides across the four replicates, depicted for the 50 most abundant *A. brevis* proteins in the haemolymph of infected ants. Proteins for which a BLAST result was found are shown in turquoise. Dashed line indicated cut-off of 15 most abundant proteins for GO prediction. B) Summary of GO term prediction of the 15 most abundant cestode proteins found in the haemolymph of infected ants with an occurrence > 3. C) Abundance of proteins that are identified as *T. nylanderi* or *A. brevis* within the haemolymph of infected ants. D) Share of *A. brevis* proteins with or without SignalP sites indicative of secretion. Shown are proteins that are found in at least 1, at least 2, at least 3 and in all biological replicates.

The mean share of cestode proteins in the haemolymph of infected ants amounted to 7% of all identified proteins across the four biological replicates (Figure 2 C). Of the 15 most abundant cestode proteins, only one (thioredoxin peroxidase) could be annotated (Figure 2 A). For the remaining candidates, we procured a prediction for functional GO terms, of which transport, regulation of metabolic process, regulation of nitrogen compound metabolic process, and cellular localization were found for all 15 proteins (Figure 2 B).

More proteins were identified from whole cestode samples (3,188) compared to cestode proteins detected in *T. nylanderi* haemolymph (263). Of those proteins, 3,035 are uniquely found in whole cestodes, whereas 110 are uniquely found in the haemolymph, possibly due to dynamic range detection by mass spectrometry and/or low abundance in whole cestode samples. The proportion of unknown proteins (i.e., proteins that had no BLASTp hit at an e-value of 1e-5) was much higher in the haemolymph samples (66.9%) compared to whole cestode samples (16.5%). Of the 51 proteins that showed a higher relative abundance in the haemolymph compared to whole cestodes, only two proteins had an annotation (Supplement 1 Figure S2), u1 small nuclear ribonucleoprotein A and *C. briggsae* CBR-HIS-71 protein.

Based on their presence in all four replicates and a positive prediction of secretion, we identified cestode proteins that could contribute to the phenotype of infection (Table 1; for proteins that occur in at least two replicates see Supplement Table S1).

**Table 1.** Cestode proteins with secretory signals and occurrences across all four replicates that potentially contribute to the parasitized phenotype

| Protein ID | Protein Name (BLAST) | Function | Citation / organism |
| --- | --- | --- | --- |
| TRINITY_DN236_c0_g2_i3 | Peptidyl-prolyl cis-trans isomerase<br>[*Echinococcus granulosus*] | Regulation of histone<br>H3-K36 trimethylation | Ahn et al., 2016<br>*C. elegans* |
| TRINITY_DN4073_c0_g2_i2 | Thioredoxin peroxidase<br>[*Echinococcus granulosus*] | Oxidative stress<br>reduction | (Wang et al., 2019)<br>*E. granulosus* |
| TRINITY_DN4612_c0_g2_i3 | Calreticulin<br>[*Echinococcus granulosus*] | Oxidative stress<br>reduction | (Lim et al., 2014)<br>*C. elegans* |
| TRINITY_DN897_c0_g1_i2 | Superoxide dismutase [Cu-Zn]<br>[*Echinococcus granulosus*] | Oxidative stress<br>reduction | (Yang et al., 2007)<br>*C. elegans* |

The two main candidates are superoxide dismutase and thioredoxin peroxidase, which both reduce oxidative stress. Markedly, both proteins were among the cestode proteins with the highest abundance in the ant’s haemolymph (Figure 2 A). Additionally, we identified tetraspanin, a protein that is commonly used as a marker for exosomes, extracellular vesicles that enable intercellular communication (Drurey & Maizels, 2021).

Of the total amount of proteins that were found in the ant’s haemolymph, 93% could be matched to the protein database of *T. nylanderi* (Stoldt et al., 2021) and amount to 1,646 proteins. Of those proteins, 57 were significantly more abundant (p < 0.05, Welch t-test and log2 fold change > 1) in infected ants, 21 proteins in comparison to uninfected nestmates (Figure 3 A) and 36 in comparison to healthy individuals (Figure 3 B). In uninfected nestmates, 66 proteins were significantly more commonly detected, 37 compared to infected and 29 compared to healthy individuals (Figure 3 C). Healthy individuals showed an overrepresentation of 20 proteins, 19 compared to infected ants and one protein compared to uninfected nestmates of infected workers.

**Fig. 3.**
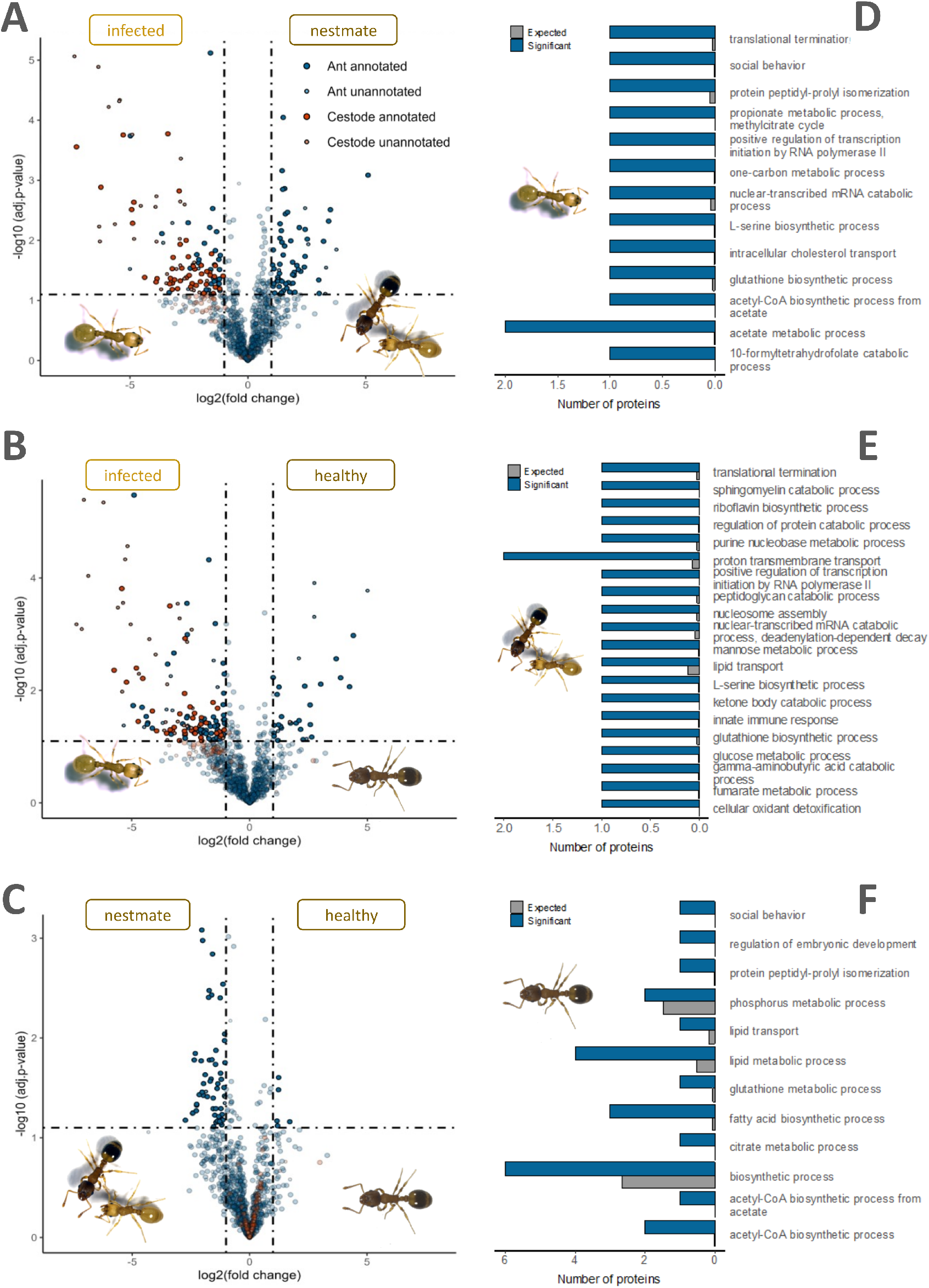
A-C) Volcano plots of pairwise comparisons between infected, nestmates of infected, and uninfected ants. Proteins, which were identified as *T. nylanderi* proteins are marked in blue and proteins, which were identified as *A. brevis* proteins are marked in red. Proteins with significantly higher abundances are depicted with stronger colours. D-F) Results of a combined GO enrichment analyses encompassing all positively enriched GO terms found in both pairwise comparisons that included D) infected ants, E) uninfected nestmates, and F) healthy ants. Shown are the number of proteins that were enriched in the specific GO functions as well as the number of proteins that were expected to be enriched, based on the number of annotated proteins.

A differential abundance analysis yielded several candidate pathways that might contribute to the observed phenotypic changes due to cestode infection (Table 2; for complete list see Supplement 2 A), many of which were postulated to extend lifespan in ants and other insects (Currin-Ross et al., 2021; Fang et al., 2014; Słowińska et al., 2019; Su et al., 2021), including sequestosome-1, chymotrypsin-1, and chymotrypsin-2, while one protein with additional functions in immunity, ras-related protein Rac1, is postulated to decrease lifespan in *Drosophila* (Slack, 2017). Infected ants showed a higher abundance of proteins that are implicated in GO functions such as cytosolic 10-formyltetrahydrofolate catabolic process, innate immune response, or L-serine biosynthetic process (Figure 3 D, for complete list see Supplement 1 Table S2, Supplement 1 Table S3).

**Table 2.** Candidate genes for facilitating the parasitised phenotype, resulting from a differential abundance analysis

| Protein ID | Protein name (BLAST) | Function | Citation / organism |
| --- | --- | --- | --- |
| MSTRG.11757.1.p1 | Death-associated protein 1<br>[*Solenopsis invicta*] | Regulator of apoptosis | Xia et al., 2017<br>*Marsupenaeus japonicus* |
| MSTRG.8546.2.p1 | Chymotrypsin-1<br>[*Temnothorax curvispinosus*] | Deubiquitination | Nguyen et al., 2019<br>*D. melanogaster* |
| MSTRG.8542.1.p1 | Chymotrypsin-2<br>[*Temnothorax curvispinosus*] | Deubiquitination | Nguyen et al., 2019<br>*D. melanogaster* |
| MSTRG.7977.1.p1 | Chymotrypsin-2<br>[*Temnothorax curvispinosus*] | Deubiquitination | Nguyen et al. 2019<br>*D. melanogaster* |
| MSTRG.8184.1.p1 | Sequestosome-1<br>[*Temnothorax curvispinosus*] | Autophagy, inactivation of<br>TOR | Negróni et al., 2021<br>*T. rugatulus* |
| MSTRG.174.1.p1 | Vitellogenin-like A<br>[*Temnothorax curvispinosus*] | Division of labour, brood<br>care | Kohlmeier et al., 2018<br>*T. longispinosus* |

We also noticed an overabundance of the protein very long-chain-fatty-acid–CoA ligase bubblegum, which has recently been found to increase in expression with age in termite queens (Séité et al., 2022). Finally, infected individuals compared to nestmates and healthy individuals, showed a seven- and 18-times higher abundance of Vg-like A (Supplement 1 Figures S4-5), a protein that elicits brood care behaviour in *Temnothorax longispinosus* (Kohlmeier et al., 2018) and *Diacamma sp*. workers (Miyazaki et al., 2021).

In their non-infected nestmates, we found a higher abundance of proteins that reduce lifespan in other species, such as calpain-A, ras-related protein Rac1, and beta-1,3-glucan-binding protein (through upregulation of toll-signalling pathway), as well as peroxiredoxin-6, which has a positive influence on lifespan through detoxification (Quigley et al., 2018). For the results of a GO term enrichment analysis in non-infected nestmates and healthy individuals, see Supplement (Supplement 1 Tables S4-7).

We compared the proteome of queens to the proteome of healthy and infected workers in a differential abundance analysis to find proteins that long-lived infected workers have in common with long-lived queens. Two proteins were more abundant in queens compared to healthy workers, icarapin and Vg-like-A (Supplement 1 Figure S3). Those two proteins were also more abundant in infected workers compared to both healthy workers (Supplement 1 Figure S4) and nestmate workers (Supplement 1 Figure S5).

We furthermore checked for proteins that are uniquely detected in both healthy queens and infected workers and found fatty acid-binding protein 1, liver-like and facilitated trehalose transporter Tret1-like (Supplement 1 Figure S6). The latter protein regulates the level of trehalose within the haemolymph (Elbein et al., 2003). In healthy queens we found 30 different proteins that were unique to them as compared to healthy workers (Supplement 1 Table S8 and Figure S7). Of those, 24 proteins were uniquely shared with infected workers (Supplement 1 Table S8), of which four were previously found in the enrichment analysis in infected workers (two uncharacterized proteins, death-associated protein 1, and chymotrypsin-2). While conclusive studies on its function in insects are missing, death-associated protein 1 positively regulates apoptosis in shrimp (Xia et al., 2017), while chymotrypsin-2 was shown to extend lifespan in *Drosophila* (Nguyen et al., 2019).

## Discussion

Parasites have found many fascinating ways to adapt to their hosts. These adaptations are largely aimed at exploiting host resources, evading immune responses and increasing their chances of transmission to the next host. The latter category in particular has been identified as a target for parasite-mediated host manipulation. Cases, in which parasites actively alter the behaviour of their hosts to facilitate the encounter between intermediate and final host, are well known (Berdoy et al., 2000; Golcu et al., 2014; Lacroix et al., 2005; Thomas et al., 2002). But not only the potentially targeted tissues or pathways can be diverse, also the mechanisms, by which parasites can influence them, are. Parasites can cause changes in the host, for example, by their location alone, by energy drain, by inflammation of neural tissue, by influencing the host’s epigenome, or by protein-protein interactions (Doherty & Matthews, 2022; Lafferty & Shaw, 2013; Martin et al., 2015). By analysing the parasitic proteins that can be found in the host’s haemolymph, we gain insight into the latter mechanism. We however not only analysed proteins with parasitic origin, but also the host’s proteome and found significant changes when comparing infected to healthy individuals. Those changes could be products of epigenetic manipulation and thus signs for adaptive manipulation. They could however also be signals of untargeted, but nevertheless beneficial, by-products of the infection. In general, active manipulation is particularly difficult to prove because the symptoms of infection and exploitation are complex and their ultimate causes are difficult to separate from untargeted side effects, such as the host’s immune response (Heil, 2016). We therefore urge for caution when interpreting the results of the ant proteome, and will discuss them accordingly in later sections of the discussion.

### Parasite proteins in the host haemolymph

Here we were particularly interested in determining the molecular basis of the extended lifespan of infected workers. In social insects, the life expectancy of individuals belonging to different castes can vary vastly. Reproductive individuals generally live much longer and show little sign of senescence compared to their worker counterparts. Oxidative stress is often discussed as the main cause of senescence (Finkel & Holbrook, 2000), but the links between oxidative stress, antioxidant processes and lifespan seem to be highly species-specific in social insects (de Verges & Nehring, 2016; Kramer et al., 2021; Lucas & Keller, 2014; Majoe et al., 2021). According to the oxidative stress theory of ageing (Finkel & Holbrook, 2000) long-lived organisms should be characterised by enhanced molecular repair capabilities, lower rates of molecular damage (i.e., lower production of reactive oxygen species or fewer replication defects) and/or higher antioxidant production. Manipulation of any of these mechanisms could alter the ability of individuals to cope with oxidative stress and thus influence the progression of senescence. Indeed, among the most highly secreted (annotated) cestode proteins, we mainly found proteins with a potential function in oxidation reduction. However, infection adds another dimension to this picture, as oxidation is a common strategy of an insect’s immune system in the defence against parasites, especially by using encapsulation and melanisation (reviewed in Zug & Hammerstein, 2015). In turn, parasites release antioxidants to counter those attacks and ensure a successful infection (de Bekker et al., 2015; Mesías et al., 2019; Piacenza et al., 2013). We would nevertheless argue that the most plausible reason for the release of such antioxidants is to actively prolong the lifespan of the ant host, as we did not find an enrichment of ant proteins with oxidative functions, and considering that we only analysed adult ants with a (potentially long) established infection, which takes place during the larval stage, making it unlikely that the release of antioxidants is targeted at countering an attack by the host.

For most of the proteins released by the cestode into the ant’s haemolymph, no functional annotations could be found. This stands in striking contrast to the complete proteome of the cestode, where more than 80% of the proteins could be annotated. The closest relative with an annotated genome, *Echinococcus granulosus* (Korhonen et al., 2022), belongs to a different family, the *Taeniidae*; thus, it was to be expected to find proteins that are not present in the other species. However, the profound discrepancy in the share of novel proteins between the secretome and proteome of the cysticercoid was surprising in its extent and might reflect unique adaptations specific to this parasite-host interaction. Indeed, similar cases have been presented in fungal and cestode parasites, where the authors found many novel proteins that the parasites release during infection (Berger et al., 2021; de Bekker et al., 2015). Functional annotations, especially of the most secreted proteins, could reveal new and specific properties important for infection dynamics or lifespan extension.

One of the most important challenges to overcome for any parasite is the host’s immune system. Many protozoan or helminthic parasites chose immune-privileged sites (i.e., sites that are less targeted by immune responses), employ antigenic variation or masking, molecular mimicry, antibody trapping, or secrete proteases to evade the immune system (Cortés et al., 2017; Flisser et al., 1986). Dissections of the infected ants so far showed no detectable immune response in the form of encapsulation or melanisation (Stoldt et al., 2021), which indicates successful evasion of the immune system, although the mechanism used is so far unknown. One of the most abundant cestode proteins that we found in the haemolymph of infected ants, acylneuraminate cytidylyltransferase, might provide a possible explanation for this. This protein synthesizes CMP-Neu5Ac, the activated form of N-Acetylneuraminic, which constitutes the basic monomer of polysialic acid (Bravo et al., 2001). Polysialic acid, either as a capsule or as components of mucins, are used by different parasites and seem to convey a means to escape the host immune response through molecular mimicry (Ghosh, 2020). Parasites not only attempt to modify host behaviour or physiology through the release of proteins. Especially helminths are known to use exosomes to facilitate intercellular parasite-host communication (Coakley et al., 2016). Exosomes can contain proteins, but also lipids and nucleic acids such as miRNAs (Coakley et al., 2016; Drurey & Maizels, 2021). Especially the latter are proposed to interfere with host gene expression and thus directly influence the production of proteins (reviewed in Britton et al., 2020). We found tetraspanin, which is a known marker for exosome activity, in all replicates of cestode proteins within the ant haemolymph, suggesting that transfer of materials occurs between the cells of host and cestode. In line with this, a gene expression study on the same system found vesicle-mediated transport to be enriched in cestodes from ants with low parasitic load compared to highly infected individuals (Sistermans et al. unpublished).

### Infection-induced changes of the ant proteome

Infected ants exhibit a mixture of traits characteristic for young workers (morphology, high fat content, metabolic rate; Beros et al., 2021) and for queens (increased longevity, reduced activity, high reproductive potential; Beros et al., 2019). Since caste differences in social insects are usually not due to genetic differences, but controlled by differential gene expression, hijacking pre-existing regulatory pathways that make an individual more queen-like might be an elegant strategy from the parasite’s point of view. However, as discussed earlier, proving those mechanisms experimentally is nearly impossible, as those changes in phenotype might be by-products of the infection, and as such, although beneficial to the parasite, cannot be regarded as adaptive host manipulation. We nevertheless would like to highlight potential pathways and genes that constitute potential targets of adaptive manipulation according to our findings. One objective was to find proteins that might facilitate the more queen-like traits of the infected workers, i.e., proteins that queens and infected workers have in common compared to healthy workers. A previous transcriptome study on this system found no more overlap in gene expression between queens and infected workers than would be expected by chance, albeit both overexpress carboxypeptidase B, which in *D. melanogaster* is encoded by the silver gene and linked to prolonged lifespan (Carnes et al., 2015; Stoldt et al., 2021). Due to posttranscriptional and post-translational regulation mechanisms, mRNA expression will not directly translate 1:1 to protein abundances (Gunawardana & Niranjan, 2013) and might thus only show parts of the picture. Proteomic data can thus help to improve our understanding how an infection translates into an infection phenotype, and furthermore allows us to sample and analyse processes that are happening in the haemolymph. Indeed, we found two candidates that were more abundant in both groups. One of them is vitellogenin-like A. In social hymenopterans, Vg genes have diversified and are especially known for their functions with regards to division of labour (Corona et al., 2013; Kohlmeier et al., 2018; Morandin et al., 2014). The abundance of certain copies of Vg in honey bees positively correlates with worker lifespan (Corona et al., 2007; Münch & Amdam, 2010). Vg is preferentially carbonylated and thus functions as a buffer to oxidative damage for other proteins (de Verges & Nehring, 2016), which facilitates higher oxidative stress resistance in the honey bee (Seehuus et al., 2006). We found higher Vg-like A abundance in infected individuals, which might contribute to protection from oxidative damage and an increased lifespan. Since Vg-like A specifically has been shown to control brood care behaviour in *Temnothorax* ants (Kohlmeier et al., 2018), a higher abundance of this protein might explain the preference of infected individuals to stay close to the brood (although they exhibit little brood care behaviour). The fact that this protein is more abundant in both queens and infected workers underlines its putative role in facilitating the queen-like phenotype of infected individuals regarding both behaviour and lifespan.

We furthermore found two proteins exclusively in infected workers and healthy queens: fatty acid-binding protein 1, and facilitated trehalose transporter Tret1-like. Especially the latter is another interesting candidate for further elucidation of the long-lived phenotype of the infected workers, as this protein is responsible for transporting trehalose, the main source of energy for insects, from the fat body into the haemolymph. There, one of trehalose’s functions is the protection from different environmental stressors, amongst them oxidative stress (Elbein et al., 2003), against which infected workers seem to be particularly resistant (Beros et al. unpublished data). Our study system is special in many ways, as one might argue that on an individual level, an infection is beneficial (longer lifespan, increased reproductive potential (Beros et al., 2019)), however, the colony as a whole suffers fitness losses (Beros et al., 2015; Scharf et al., 2012). Following and expanding on Dawkins’ theory of the extended phenotype (1982), in which, classically, one party influences the expressed phenotype of another party beyond the reach of their normal physical constraints, we find consequences of parasitism on the level of the whole colony. Previous studies showed that infected colonies are less aggressive and uninfected workers live shorter (Beros et al., 2015, 2021), and furthermore exhibit signs of stress by raising more queen-worker intercastes and displaying a male-biased sex ratio (Beros et al., 2021). Also on a level of gene expression, the influence of a cestode infection on uninfected nestmates could be shown (Feldmeyer et al., 2016). It is thus not surprising that we found alterations in the proteome of uninfected nestmates compared to workers from uninfected colonies. Markedly, uninfected nestmates showed an enrichment of processes that are responsible for detoxification and immune response. This could be a consequence of the observed shifts in the colonies’ general stress level (Beros et al., 2021) that could lead to a higher susceptibility to pathogens, which would explain the higher expression of those pathways.

## Conclusion

We were able to identify proteins actively secreted into the host with functions in oxidation reduction and putative immune escape properties that can (i) explain how the cestode is able to maintain host manipulation and that (ii) likely contribute to the exhibition of the observed infection phenotype of their host with a substantially prolonged lifespan. On another level, we are able to add to studies that show that not only the infected individual carries the consequences of an infection, but that the whole colony is affected. For most secreted proteins annotated orthologs could not be found, which is indicative of potential novel functions and a long history of adaptation within this species interaction. Especially for the most abundant proteins no matches in sequence similarity could be found. We found a marker for exosome activity, suggesting cell-cell communication from parasite to host. Within the host’s proteome, we identified proteins that are subject to parasite-induced shifts in abundance and that are prime candidates to explain the infection phenotype with its extended lifespan and behavioural alterations. Amongst them is vitellogenin-like A, a protein with functions in the division of labour and fecundity. Future studies should focus on the identification of unknown novel proteins, either by structural predictions or knock-outs.

## ACKNOWLEDGEMENTS

The authors wish to thank Romain Libbrecht, Joe Colgan, Marie-Pierre Meurville and Hugo Darras for their time and effort to provide insightful comments on a first version of this manuscript. This project was funded by the Deutsche Forschungsgemeinschaft (DFG, German ResearchFoundation) – GRK2526/1 – Projectnr. 407023052.

Processed label-free quantitative proteomics data are provided in Supplementary Data 4. Raw mass spectrometry data will be uploaded to a suitable platform upon publication of the manuscript.

## Notes

### Competing Interest Statement

The authors have declared no competing interest.

### Summary of Updates

Title

