## Supplement 1 for "Long live the host! Proteomic analysis reveals possible strategies for parasitic manipulation of its social host"

### Supplementary Material

**Table S1**: Cestode proteins with secretory signals and occurrences in at least two of the four replicates that potentially contribute to the infection phenotype

| Protein ID | BLAST result | Function (Uniprot *Cele*) |
|---|---|---|
| TRINITY_DN10416_c0_g1_i5 | Dynein light chain 1, cytoplasmic [*Echinococcus granulosus*] | apoptosis, Golgi - ER transport |
| TRINITY_DN10876_c0_g2_i1 | Mediator of RNA polymerase II transcription subunit 15 (Mediator complex subunit 15) [*Echinococcus granulosus*] | determination of adult life span, transcription |
| TRINITY_DN11884_c0_g1_i1 | Dynein light chain 2 [*Schistosoma mansoni*] | apoptosis |
| TRINITY_DN13270_c0_g1_i7 | Nuclear hormone receptor family member fax-1 [*Echinococcus granulosus]* | regulation of gene expression |
| TRINITY_DN1512_c0_g2_i7 | Kinesin light chain (KLC) [*Echinococcus granulosus]* | probable organelle transport |
| TRINITY_DN17541_c0_g1_i1 | Dynein light chain 1 [*Echinococcus granulosus]* | apoptosis, regulator of RNA-binding protein |
| TRINITY_DN1986_c0_g2_i1 | Lysosomal aspartic protease [*Echinococcus granulosus]* | transmembrane transport |
| TRINITY_DN2071_c0_g3_i1 | Sodium/potassium-transporting ATPase subunit beta-1 [*Echinococcus granulosus]* | transport across plasma membrane |
| TRINITY_DN21918_c0_g2_i4 | Multidrug resistance protein pgp-1 (EC 7.6.2.2) (P-glycoprotein A) (P-glycoprotein-related protein 1) [*Echinococcus granulosus*] | immune response |
| TRINITY_DN236_c0_g2_i3 | Peptidyl-prolyl cis-trans isomerase [*Echinococcus granulosus*] | regulation of histone H3-K36 trimethylation |
| TRINITY_DN2636_c0_g1_i1 | Heat shock protein [*Echinococcus granulosus*] | increase of lifespan |
| TRINITY_DN2696_c0_g2_i2 | Fatty acid-binding protein homolog 8 [*Echinococcus granulosus*] | lipid transport |
| TRINITY_DN2981_c0_g2_i1 | Calmodulin [*Echinococcus granulosus*] | increase of lifespan, regulation of gene expression |
| TRINITY_DN314_c0_g2_i4 | Calmodulin [*Caenorhabditis remanei*] | increase of lifespan, regulation of gene expression |
| TRINITY_DN3186_c0_g2_i2 | Fatty acid-binding protein [*Echinococcus granulosus*] | lipid transport |
| TRINITY_DN37546_c0_g1_i1 | Multidrug resistance protein 3 [*Echinococcus granulosus*] | immune response |
| TRINITY_DN3867_c0_g1_i2 | protein TFG [*Diabrotica virgifera virgifera*] | downregulation of apoptosis |
| TRINITY_DN3916_c0_g1_i1 | Histone H4 [*Echinococcus granulosus*] | regulation of gene expression |
| TRINITY_DN4028_c0_g2_i1 | Protein disulfide-isomerase [*Echinococcus granulosus*] | stress response |
| TRINITY_DN40716_c0_g1_i1 | nascent polypeptide-associated complex subunit alpha [*Drosophila albomicans*] | stress response |
| TRINITY_DN4073_c0_g2_i2 | Thioredoxin peroxidase [*Echinococcus granulosus*] | oxidative stress reduction |
| TRINITY_DN4276_c0_g2_i1 | Clathrin heavy chain [*Echinococcus granulosus*] | determination of adult life span |
| TRINITY_DN4612_c0_g2_i3 | Calreticulin [*Echinococcus granulosus*] | oxidative stress reduction |

| | | |
|---|---|---|
| TRINITY_DN771_c0_g1_i4 | Small heat shock protein p36 [*Echinococcus granulosus*] | increase of lifespan |
| TRINITY_DN531_c0_g2_i3 | MANF/CDNF protein [*Echinococcus granulosus*] | stress response |
| TRINITY_DN897_c0_g1_i2 | Superoxide dismutase [Cu-Zn] (EC 1.15.1.1) [*Echinococcus granulosus*] | oxidative stress reduction |
| TRINITY_DN9024_c0_g1_i1 | Sodium/hydrogen exchanger 2 [*Echinococcus granulosus*] | transport |

**Table S2**: Results of GO term enrichment analysis of proteins that are overrepresented in infected individuals in comparison to their uninfected nestmates

| GO.ID | Term | Annotated | Significant | Expected | Fisher |
|---|---|---|---|---|---|
| GO:0006083 | acetate metabolic process | 4 | 2 | 0.01 | 0.0013 |
| GO:0009258 | 10-formyltetrahydrofolate catabolic proc... | 1 | 1 | 0 | 0.0013 |
| GO:0019679 | propionate metabolic process, methylcitr... | 1 | 1 | 0 | 0.0013 |
| GO:0060261 | positive regulation of transcription ini... | 1 | 1 | 0 | 0.0013 |
| GO:0019427 | acetyl-CoA biosynthetic process from ace... | 3 | 1 | 0 | 0.004 |
| GO:0035176 | social behavior | 4 | 1 | 0.01 | 0.0053 |
| GO:0006564 | L-serine biosynthetic process | 4 | 1 | 0.01 | 0.0053 |
| GO:0006730 | one-carbon metabolic process | 6 | 1 | 0.01 | 0.008 |
| GO:0032367 | intracellular cholesterol transport | 9 | 1 | 0.01 | 0.012 |
| GO:0006750 | glutathione biosynthetic process | 15 | 1 | 0.02 | 0.0199 |
| GO:0006415 | translational termination | 20 | 1 | 0.03 | 0.0264 |
| GO:0000288 | nuclear-transcribed mRNA catabolic proce... | 29 | 1 | 0.04 | 0.0381 |

**Table S3**: Results of GO term enrichment analysis of proteins that are overrepresented in infected individuals compared to healthy workers

| GO.ID | Term | Annotated | Significant | Expected | Fisher |
|---|---|---|---|---|---|
| GO:0006508 | proteolysis | 1035 | 5 | 1.23 | 0.0057 |
| GO:0006526 | arginine biosynthetic process | 9 | 1 | 0.01 | 0.0107 |
| GO:0045087 | innate immune response | 13 | 1 | 0.02 | 0.0154 |
| GO:0005975 | carbohydrate metabolic process | 655 | 3 | 0.78 | 0.0401 |
| GO:0006665 | sphingolipid metabolic process | 38 | 1 | 0.05 | 0.0443 |
| GO:0009253 | peptidoglycan catabolic process | 39 | 1 | 0.05 | 0.0454 |

**Table S4**: Results of GO term enrichment analysis of proteins that are overrepresented in nestmates compared to infected individuals

| GO.ID | Term | Annotated | Significant | Expected | Fisher |
|---|---|---|---|---|---|
| GO:0006685 | sphingomyelin catabolic process | 6 | 1 | 0 | 0.0031 |
| GO:0045087 | innate immune response | 13 | 1 | 0.01 | 0.0067 |
| GO:0006006 | glucose metabolic process | 19 | 1 | 0.01 | 0.0098 |
| GO:0006013 | mannose metabolic process | 21 | 1 | 0.01 | 0.0109 |
| GO:0006144 | purine nucleobase metabolic process | 34 | 1 | 0.02 | 0.0176 |

| | | | | | |
|---|---|---|---|---|---|
| GO:0009253 | peptidoglycan catabolic process | 39 | 1 | 0.02 | 0.0201 |

**Table S5**: Results of GO term enrichment analysis of proteins that are overrepresented in nestmates compared to healthy individuals

| GO.ID | Term | Annotated | Significant | Expected | Fisher |
|---|---|---|---|---|---|
| GO:0009231 | riboflavin biosynthetic process | 1 | 1 | 0 | 0.0013 |
| GO:0060261 | positive regulation of transcription ini... | 1 | 1 | 0 | 0.0013 |
| GO:1902600 | proton transmembrane transport | 53 | 2 | 0.07 | 0.002 |
| GO:0098869 | cellular oxidant detoxification | 2 | 1 | 0 | 0.0025 |
| GO:0046952 | ketone body catabolic process | 3 | 1 | 0 | 0.0038 |
| GO:0006106 | fumarate metabolic process | 4 | 1 | 0.01 | 0.005 |
| GO:0006564 | L-serine biosynthetic process | 4 | 1 | 0.01 | 0.005 |
| GO:0009450 | gamma-aminobutyric acid catabolic proces... | 5 | 1 | 0.01 | 0.0063 |
| GO:0042176 | regulation of protein catabolic process | 10 | 1 | 0.01 | 0.0126 |
| GO:0006334 | nucleosome assembly | 14 | 1 | 0.02 | 0.0176 |
| GO:0006750 | glutathione biosynthetic process | 15 | 1 | 0.02 | 0.0188 |
| GO:0006415 | translational termination | 20 | 1 | 0.03 | 0.025 |
| GO:0000288 | nuclear-transcribed mRNA catabolic proce... | 29 | 1 | 0.04 | 0.0361 |

**Table S6:** Results of GO term enrichment analysis of proteins that are overrepresented in healthy individuals compared to infected individuals

| GO.ID | Term | Annotated | Significant | Expected | Fisher |
|---|---|---|---|---|---|
| GO:0006633 | fatty acid biosynthetic process | 68 | 3 | 0.06 | 0.00002 |
| GO:0019427 | acetyl-CoA biosynthetic process from ace... | 3 | 1 | 0 | 0.0025 |
| GO:0006101 | citrate metabolic process | 3 | 1 | 0 | 0.0025 |
| GO:0035176 | social behavior | 4 | 1 | 0 | 0.0033 |
| GO:0006085 | acetyl-CoA biosynthetic process | 10 | 2 | 0.01 | 0.0052 |
| GO:0000413 | protein peptidyl-prolyl isomerization | 40 | 1 | 0.03 | 0.0322 |

**Table S7:** Results of GO term enrichment analysis of proteins that are overrepresented in healthy individuals compared to nestmates

| GO.ID | Term | Annotated | Significant | Expected | Fisher |
|---|---|---|---|---|---|
| GO:0045995 | regulation of embryonic development | 2 | 1 | 0 | 0.00015 |

**Table S8:** List of proteins that are unique to healthy queens as compared to healthy workers. Proteins that are uniquely shared between healthy queens and infected workers (both in comparison to healthy workers) are marked in **bold**.

| Protein ID | BLAST Results | Function (Uniprot *Dmel*) |
|---|---|---|
| MSTRG.1056.2.p1 | methylthioribulose-1-phosphate dehydratase [*Wasmannia auropunctata*] | Catalyzes the dehydration of methylthioribulose-1-phosphate |
| MSTRG.10871.1.p1 | 26S proteasome regulatory subunit 10B isoform X1 [*Ooceraea biroi*] | required for female reproduction in planthoppers (https://doi.org/10.1098/rsob.200251) |

| | | |
|---|---|---|
| MSTRG.11006.1.p1 | plastin-2 isoform X2 [*Temnothorax curvispinosus*] | NA |
| MSTRG.1166.1.p1 | eukaryotic translation initiation factor 4E transporter-like isoform X3 [*Temnothorax curvispinosus*] | Acts as a regulator of lifespan in response to cold (10.1073/pnas.1618994114), or dietary restrictions (10.1016/j.cell.2009.07.034) |
| **MSTRG.11757.1.p1** | Death-associated protein 1 [*Solenopsis invicta*] | Anti-apoptotic protein |
| **MSTRG.11759.1.p1** | Uncharacterized protein [*Temnothorax curvispinosus*] | NA |
| **MSTRG.14136.1.p1** | Tetraspanin-2A [*Temnothorax curvispinosus*] | Required for assembly of smooth septate junctions |
| **MSTRG.14279.1.p1** | V-type proton ATPase 116 kDa subunit a-like [*Temnothorax curvispinosus*] | NA |
| **MSTRG.1795.1.p1** | Uncharacterized protein [*Temnothorax curvispinosus*] | NA |
| **MSTRG.2586.9.p1** | Uncharacterized family 31 glucosidase KIAA1161 isoform X1 [*Vollenhovia emeryi*] | NA |
| **MSTRG.3123.3.p1** | Uncharacterized protein [*Contarinia nasturtii*] | NA |
| **MSTRG.3251.2.p1** | Uncharacterized protein [*Temnothorax curvispinosus*] | NA |
| **MSTRG.3401.1.p1** | ATP-dependent RNA helicase vasa isoform X2 [*Temnothorax curvispinosus*] | Involved in translational control mechanisms operating in early stages of oogenesis. |
| **MSTRG.3791.3.p1** | Aminopeptidase W07G4.4 [*Temnothorax curvispinosus*] | NA |
| MSTRG.3966.2.p1 | glutenin, high molecular weight subunit PW212 [*Monomorium pharaonis*] | NA |
| **MSTRG.4179.1.p3** | Fatty acid-binding protein 1, liver-like [*Temnothorax curvispinosus*] | Constitutes the major component of lipophorin, which mediates transport for various types of lipids in hemolymph. |
| **MSTRG.48.1.p1** | Uncharacterized protein [*Temnothorax curvispinosus*] | NA |
| **MSTRG.49.1.p1** | Uncharacterized protein [*Temnothorax curvispinosus*] | NA |
| **MSTRG.5155.1.p1** | Uncharacterized protein [*Temnothorax curvispinosus*] | NA |
| **MSTRG.5982.18.p1** | Facilitated trehalose transporter Tret1-like isoform X3 [*Temnothorax curvispinosus*] | Transport of trehalose synthesized in the fat body, thereby regulating trehalose levels in the hemolymph. |
| **MSTRG.6125.4.p1** | Carnitine O-acetyltransferase-like isoform X1 [*Temnothorax curvispinosus*] | Catalyzes the reversible synthesis of acetylcholine (ACh) from acetyl CoA and choline at cholinergic synapses. |
| **MSTRG.7157.2.p1** | Transferrin-like [*Temnothorax curvispinosus*] | Proton-conducting pore forming subunit of the membrane integral V0 complex of vacuolar ATPase. |
| **MSTRG.7742.123.p1** | N66 matrix protein-like, partial [*Temnothorax curvispinosus*] | NA |
| **MSTRG.7843.1.p1** | Slit homolog 1 protein-like [*Temnothorax curvispinosus*] | Arm and Abl proteins function cooperatively at adherens junctions in both the CNS and epidermis; critical for embryonic epithelial morphogenesis regulating cell shape changes and cell migration. |
| **MSTRG.7977.1.p1** | Chymotrypsin-2-like isoform X2 [*Temnothorax curvispinosus*] | NA |
| MSTRG.8296.1.p1 | transcriptional activator protein Pur-beta [*Temnothorax curvispinosus*] | NA |

| | | |
|---|---|---|
| **MSTRG.8544.1.p1** | Chymotrypsin-2-like [*Temnothorax curvispinosus*] | NA |
| **MSTRG.9409.3.p2** | Aminopeptidase-2 [*Temnothorax curvispinosus*] | NA |
| **MSTRG.9409.3.p5** | Aminopeptidase-2 [*Temnothorax curvispinosus*] | NA |
| **MSTRG.9616.3.p1** | Fragile X mental retardation syndrome-related protein 1 isoform X5 [*Wasmannia auropunctata*] | Polyribosome-associated RNA-binding protein that plays a role in neuronal development and synaptic plasticity |

**Table S9:** Search terms associated with longevity, stress, gene regulation and transport

| **Longevity** | **Stress** | **Gene regulation** | **Transport** |
|---|---|---|---|
| aging<br>senescence<br>lifespan<br>repair<br>piwi<br>JNK<br>apopto*<br>autophagy<br>cell death<br>oxidative<br>redox<br>TOR<br>Toll<br>toll<br>UV damage | stress | transcription factor<br>histone<br>methylation | transport<br>vesicle<br>transmembrane<br>export<br>secretion<br>excretion |

### FIGURES:

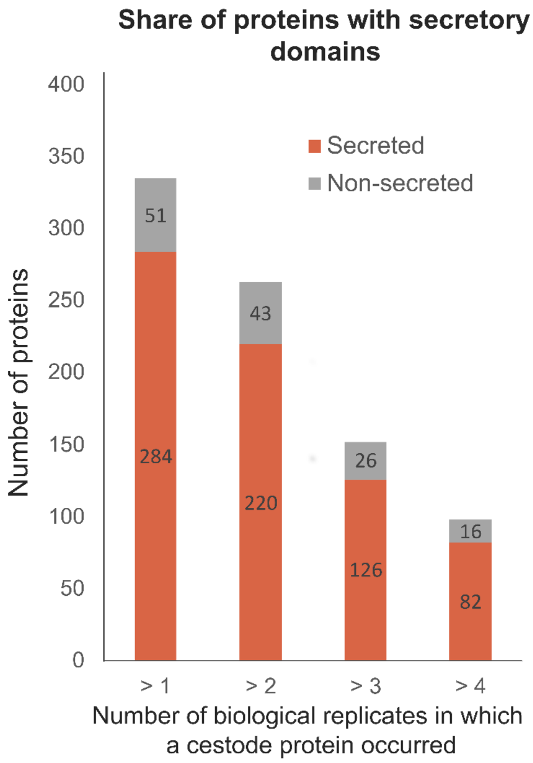


**Figure S1:** Comparison of the number of proteins that are putatively secreted (i.e., that have either a SignalP site or were marked as non-classically secreted (neural network score > 0.6) by SecretomeP across differing number of replicates that the proteins were found in.

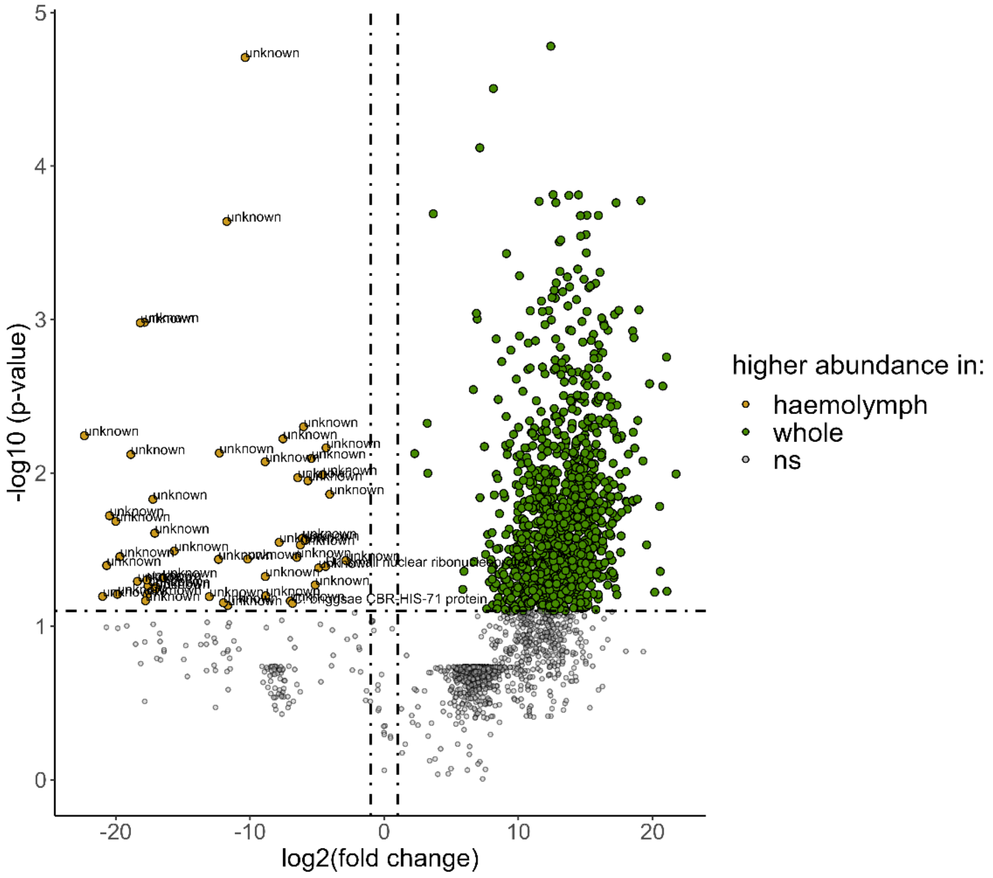


**Figure S2:** Differential abundance volcano plot showing relative protein abundances comparing cestode proteins originating from whole cestode samples and *T. nylanderi* haemolymph. Proteins with significantly different abundances are marked in either green (higher abundance in whole cestodes) or yellow (higher abundance in haemolymph). Significance level cut-offs are indicated with dashed lines. Names are only provided for proteins with higher abundances in the haemolymph.

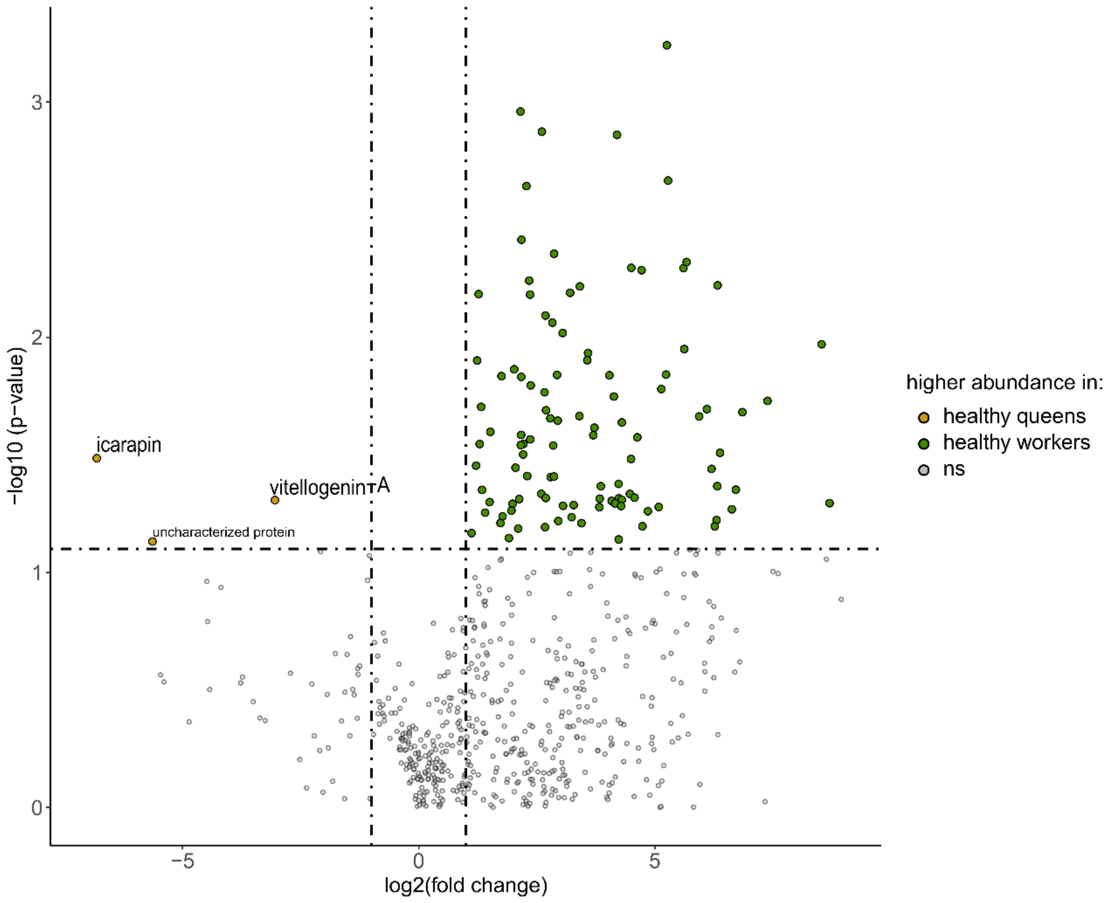


**Figure S3:** Differential abundance volcano plot showing relative protein abundances comparing healthy queens and healthy workers. Proteins with significantly different abundances are marked in either green (higher abundance in healthy workers) or yellow (higher abundance in healthy queens). Significance level cut-offs are indicated with dashed lines. Names are only provided for proteins with higher abundances in healthy queens. Proteins that are found overlapping between queens and infected workers are indicated with a larger font.

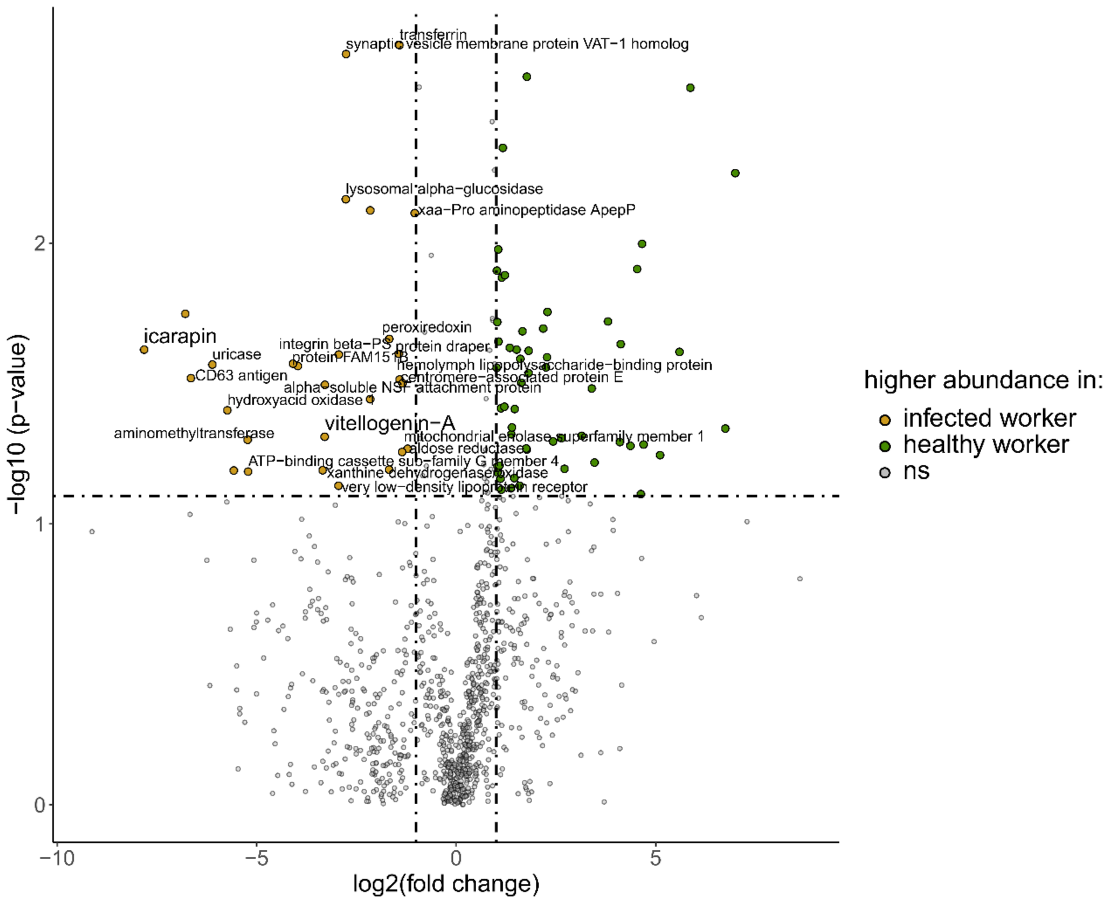


**Figure S4:** Differential abundance volcano plot showing relative protein abundances comparing infected and healthy workers. Proteins with significantly different abundances are marked in either green (higher abundance in healthy workers) or yellow (higher abundance in infected workers). Significance level cut-offs are indicated with dashed lines. Names are only provided for proteins with higher abundances in infected workers. Proteins that are found overlapping between queens and infected workers are indicated with a larger font.

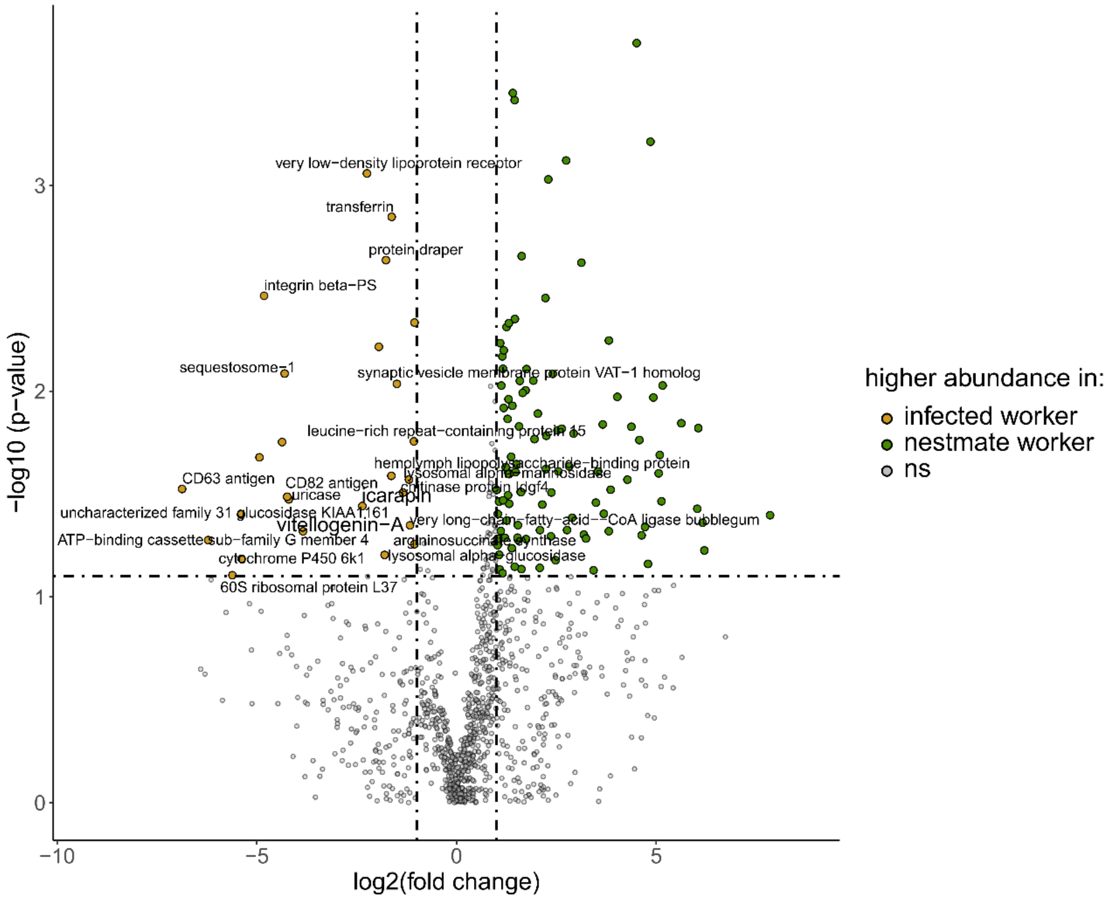


**Figure S5:** Differential abundance volcano plot showing relative protein abundances comparing infected workers and their nestmates. Proteins with significantly different abundances are marked in either green (higher abundance in nestmate workers) or yellow (higher abundance in infected workers). Significance level cut-offs are indicated with dashed lines. Names are only provided for proteins with higher abundances in infected workers. Proteins that are found overlapping between queens and infected workers are indicated with a larger font.

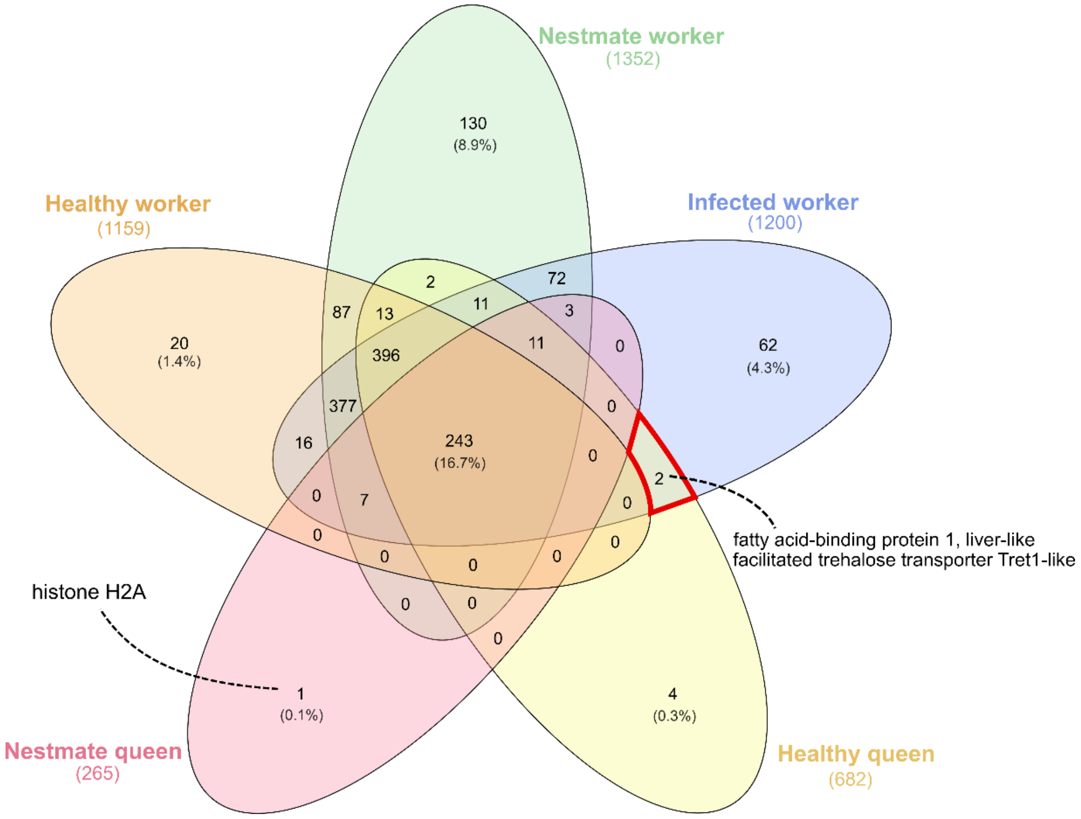


**Figure S6:** Venn diagram showing the overlap of identified *T. nylanderi* proteins between all haemolymph datasets. Only proteins that occur in at least two replicates of the respective datasets are considered. Candidate proteins that only occur in infected individuals and queens are indicated in red and their description is given in the plot. A protein that was only identified in nestmate queens (in at least 2 replicates) is Histone H2A.

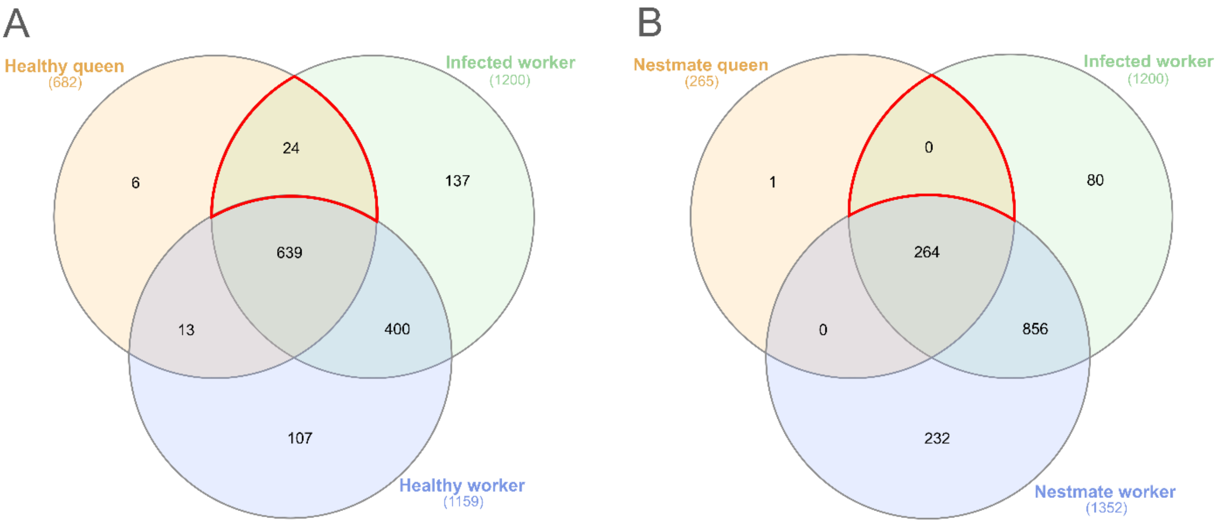


**Figure S7:** Venn diagrams showing the overlap of *T. nylanderi* proteins between different haemolymph datasets. Only proteins that occur in at least two replicates of the respective datasets are considered. **A)** comparison aimed at finding "queen-like" proteins within infected workers compared to healthy workers. **B)** comparison aimed at finding "queen-like" proteins within infected workers compared to uninfected nestmates. Candidate proteins are indicated in red.

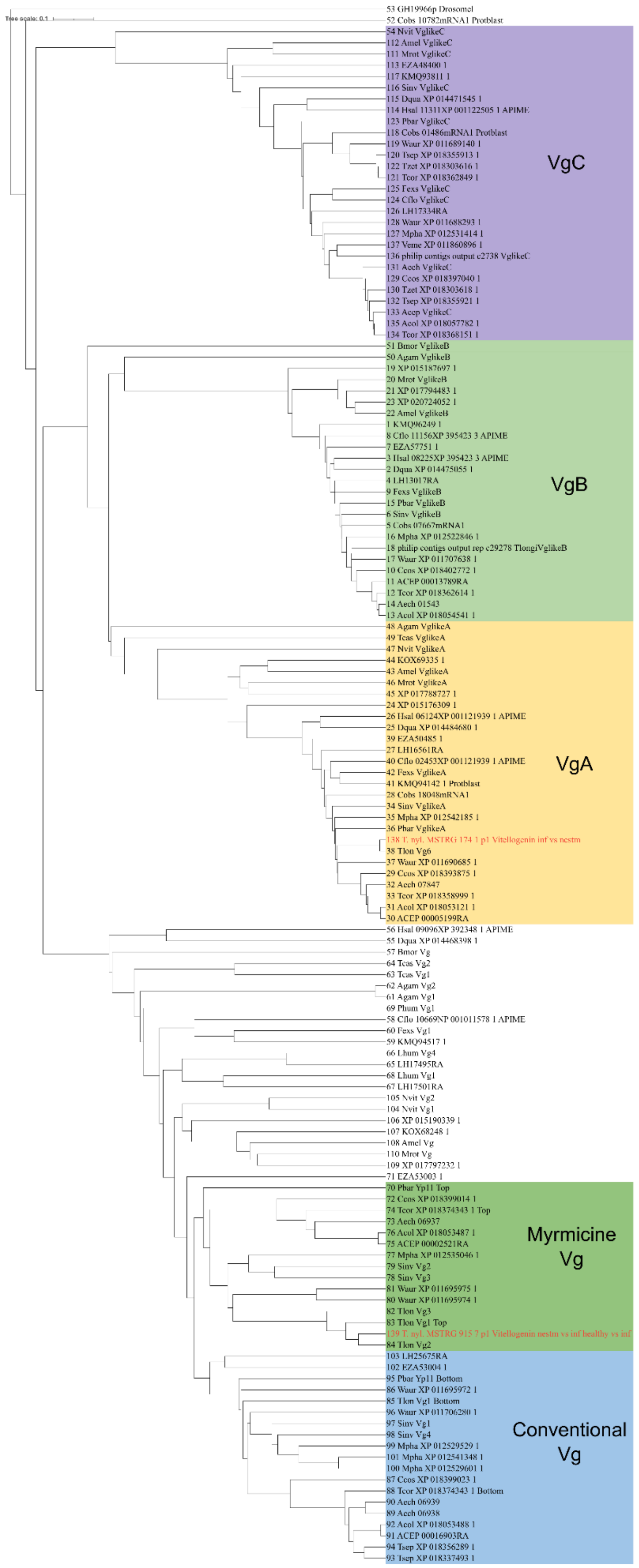


**Figure S8**: Phylogenetic tree with known vitellogenin genes. Sequences originated from Kohlmeier et al. (2016). Alignment and tree computated with MAFFT (standard settings). *T. nylanderi* sequences are highlighted in red and results of the enrichment analysis are indicated within the sequence name.
